# Machine learning-based estimation of spatial gene expression pattern during ESC-derived retinal organoid development

**DOI:** 10.1101/2023.03.26.534261

**Authors:** Yuki Fujimura, Itsuki Sakai, Itsuki Shioka, Nozomu Takata, Atsushi Hashimoto, Takuya Funatomi, Satoru Okuda

## Abstract

Organoids, which can reproduce the complex tissue structures found in embryos, are revolutionizing basic research and regenerative medicine. In order to use organoids for research and medicine, it is necessary to assess the composition and arrangement of cell types within the organoid, i.e., spatial gene expression. However, current methods are invasive and require gene editing and immunostaining. In this study, we developed a non-invasive estimation method of spatial gene expression patterns using machine learning. A deep learning model was trained with an encoder-decoder architecture on a dataset of retinal organoids derived from mouse embryonic stem cells. This method successfully estimated spatially plausible fluorescent patterns with appropriate intensities, enabling the non-invasive, quantitative estimation of spatial gene expression patterns within each tissue. Thus, this method could lead to new avenues for evaluating spatial gene expression patterns across a wide range of biology and medicine fields.

**Highlights:**

- A non-invasive estimation method of spatial gene expression pattern is proposed
- A CNN architecture is employed to convert a phase-contrast to fluorescence image
- The method was trained on a dataset of mouse ESC-derived retinal organoids
- Spatially plausible patterns of Rx gene expressions were successfully estimated

## Introduction

Stem cell-based tissue engineering has become essential for understanding organogenesis and has gained more attention in regenerative medicine (Grassi et al., 2019; Nakamura and Sato, 2018; Nath et al., 2022; Perez-Gonzalez et al., 2022; Sasai, 2013; Takebe and Wells, 2019; Zhao et al., 2022). For example, the loss of our body parts due to injury, aging, or disease can lead to serious life-threatening conditions, which could be slowed down or reduced by cell transplantation therapy (Madrid et al., 2021; Mandai, 2023). However, autologous cell therapy using induced pluripotent stem (iPS) cells, established from patients, requires purification of cell lines and generation of high-quality target cells to avoid unwanted cell differentiation or contamination. This requirement is even more important for the transplantation of stem-cell derived tissues and organs, which require a well-structured complex of different cell types.

Recently, *in vitro* models derived from pluripotent stem cells, referred to as organoid, are useful for better understanding organogenesis. We have also developed several models of neuroepithelia, such as mouse embryonic stem cell (ESC)-derived retinal organoids (Eiraku et al., 2011; Hasegawa et al., 2016; Okuda et al., 2018; Takata et al., 2017). These organoids recapitulate three-dimensional complex structures and developmental processes as observed in organogenesis. Organoids typically contain more cell types than tissues obtained in conventional planar differentiation culture (Kanton et al., 2019; Lancaster et al., 2013). Therefore, organoid technologies have the potential to serve as a source of cells, tissues, and organs for transplantation and as a screening platform for drug discovery via patient-relevant cell sources (Beghin et al., 2022; Driehuis et al., 2020; Schuster et al., 2020).

Grown organoids are evaluated by the expression of a master gene, such as Rx (aka Rax) for eye field, since the level of gene expression determines cell fate, and its location and timing correspond to those of tissue differentiation. To monitor the genetic activity of cells within each organoid, fluorescence-based reporters have been used along with homologous recombination-based knock-in and genome editing tools (Takata and Eiraku, 2018) (Artegiani et al., 2020 from other groups that use fluorescent-based reporter systems). These reporters allow us to reveal the location and timing of target gene expression during the formation of each organoid, in which different cell types arise in a stepwise manner. Moreover, since the fluorescence intensity of the reporter reflects the level of gene expression, it is also possible to quantitatively determine the spatiotemporal distribution of gene expression levels.

Although genetic analyses have revealed important genes and regulatory genomic regions, genomic alterations require the inactivation of at least one of two endogenous alleles. Therefore, a complete understanding of endogenous gene networks is missing and ethical issues preclude the use of genomic alterations for cell, tissue, and organ transplantation. Moreover, genomic alterations require substantial time and effort. The current remaining method is to manually select target regions in individual samples from brightfield or phase-contrast images, however this method has limitations of throughput and subjective classification criteria that can lead to high variability among observers. In addition to the above issues, ethical issues preclude the use of genomic alterations for cell, tissue, or organ transplantation into humans. To overcome these limitations, we developed a new method to automatically and non-invasively estimate spatial patterns of specific gene expressions in each organoid using machine learning.

Machine-learning approaches have been applied to stem cell biology, especially, isolated cells differentiated from stem cells on planar dishes (Coronnello and Francipane, 2022; Waisman et al., 2019; Zhu et al., 2021). In these studies, deep learning models were trained on a paired dataset of either brightfield or phase-contrast images and fluorescent images obtained by immunostaining. These models were able to classify individual cell types based on either brightfield or phase-contrast images.

Furthermore, machine learning approaches are beginning to be applied to organoids. A pioneering work applied them to retinal organoids and succeeded in predicting whether each organoid differentiate into retina or not based on brightfield images (Kegeles et al., 2020). However, estimating spatial and temporal gene expression remains challenging.

In this study, we developed a machine learning-based technique to estimate the spatiotemporal distribution of gene expression levels within each organoid, based on a phase-contrast image. We formulate the problem as an image transfer, i.e., a convolutional neural network (CNN) with an encoder-decoder architecture takes a phase-contrast image as input and directly outputs a fluorescent image with a spatial distribution of gene expression (Fig. 1A). Differing from a previous method that simply classifies each tissue as either differentiated or not, our method estimates pixel-wise intensities within each tissue. Experiments demonstrated that our method succeeded in estimating spatially plausible fluorescent patterns with appropriate intensities.

**Figure 1.**
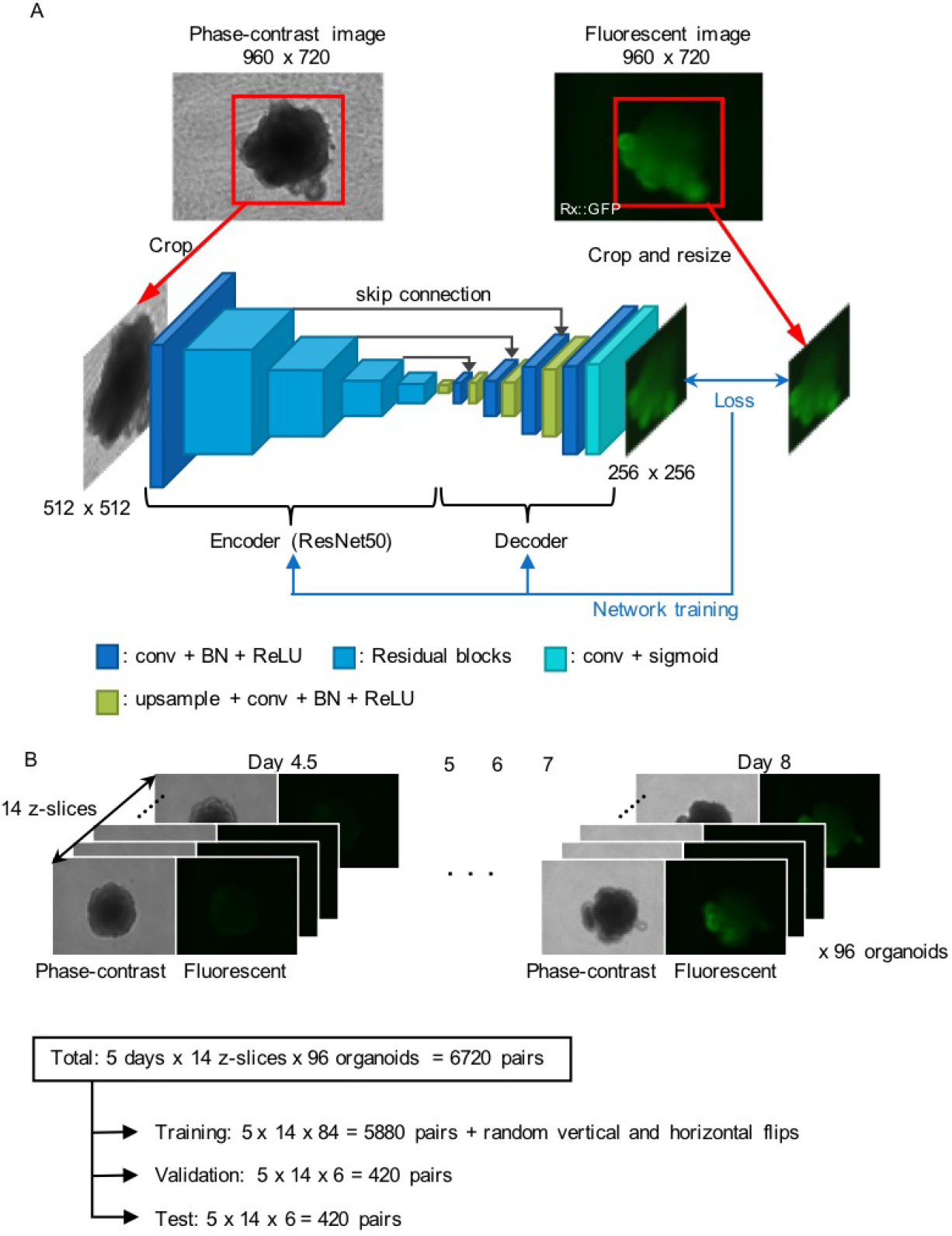
Network architecture and dataset for estimating spatial gene expression patterns. (A) Network architecture. Our network has an encoder-decoder architecture, which takes a phase-contrast image as input and then outputs a fluorescent image. The encoder backbone is the ResNet50 with four residual blocks, and the decoder has upsampling layers to increase the resolution of the intermediate feature map. (B) Our dataset consists of pairs of phase-contrast and fluorescent images of mouse ESC-derived retinal organoids. The fluorescence is of a GFP, which gene was knocked in under the promoter of a master gene of retinal differentiation, Rx. The dataset was divided into the training, test, and validation subsets.

## Results

### CNN architecture and dataset for estimating spatial gene expression patterns

Our model utilizes a CNN that takes a phase-contrast image as input and estimates a fluorescent image as output (Fig. 1A). The typical input to a CNN is a two-dimensional (2D) image. This 2D image is passed through several convolution layers, each followed by a nonlinear activation function. The training parameters correspond to the weights of these convolution kernels and the biases. Our network has a U-Net-like architecture (Ronneberger et al., 2015), which is an encoder-decoder structure with skip connections. The embedded features from the encoder are passed through the decoder, which consists of upsampling and convolution layers to increase the resolution of the intermediate feature maps to obtain a fluorescent image as output.

In our model, the ResNet50 (He et al., 2016) was used as the backbone of the encoder. The size of the input image for the ResNet50 is 3 × *H* × *W*. To use the pre-trained model of the ResNet50, gray-scale phase-contrast images were replicated in the axis of the channel to create three-channel images. At the first layer, a convolution with stride 2 is applied to the input image to generate features of size 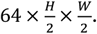. The ResNet50 has 4 residual blocks and the size of the output features of these blocks are 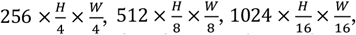, and 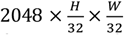, respectively. These features are then concatenated to the decoder to exploit multi-scale information. The output of the decoder is a fluorescent image of size 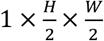. Note that each convolution layer has a batch normalization (BN) layer and a rectified linear unit (ReLU) activation function, except for the final convolution layer, which has a sigmoid activation function to constrain the range of the output values between 0 and 1.

The network was optimized by minimizing the training loss computed on the output and corresponding ground-truth fluorescent images. The combination of mean squared error (MSE) and cosine similarity, which captures structural patterns from the entire image, was used as the training loss.

To train, validate, and test our model, we cultured retinal organoids derived from mouse ESCs using the SFEBq method (Eiraku et al., 2011). In this culture, a GFP gene was knocked-in under the promoter of a master gene of retinal differentiation, Rx. Using this method, we obtained a dataset of a pair of phase-contrast image and fluorescent image of Rx during retinal differentiation (Fig. 1B). Images were captured for 96 organoids at 4.5, 5, 6, 7, and 8 days after the start of SFEBq, where each sample was captured as 14 Z-stack images. This resulted in a total of 96 × 5 × 14 = 6720 image pairs were obtained. These image pairs were divided into 5880, 420, and 420 samples for training, validation, and test, respectively. For data augmentation, we randomly flipped the input images vertically and horizontally during training. While the image resolution of both phase-contrast and fluorescent images is 960 × 720, the 512 × 512 regions where organoids appear were extracted.

### Machine learning successfully estimates spatial Rx expression patterns

To demonstrate our model, we applied it to 420 samples of the test data. As a result, the proposed model successfully estimated the spatial expression patterns of Rx from phase-contrast images during retinal organoid development (Fig. 2). During development, multiple optic vesicles are formed through large and complicated deformations (Fig. 2A). This process begins with a spherical embryonic body, with some portions of the tissue surface evaginating outward to form hemispherical vesicles, i.e., optic vesicles. Importantly, the resulting morphology of retinal organoids, especially optic vesicles, varies widely (Decembrini et al., 2014). This process is known to be governed by the expression of the Rx gene (Fig. 2B). That is, the Rx gene is gradually expressed in several parts of the tissue surface, so-called eye field, where cells differentiate from neuroepithelium into several types of retinal cells.

**Figure 2.**
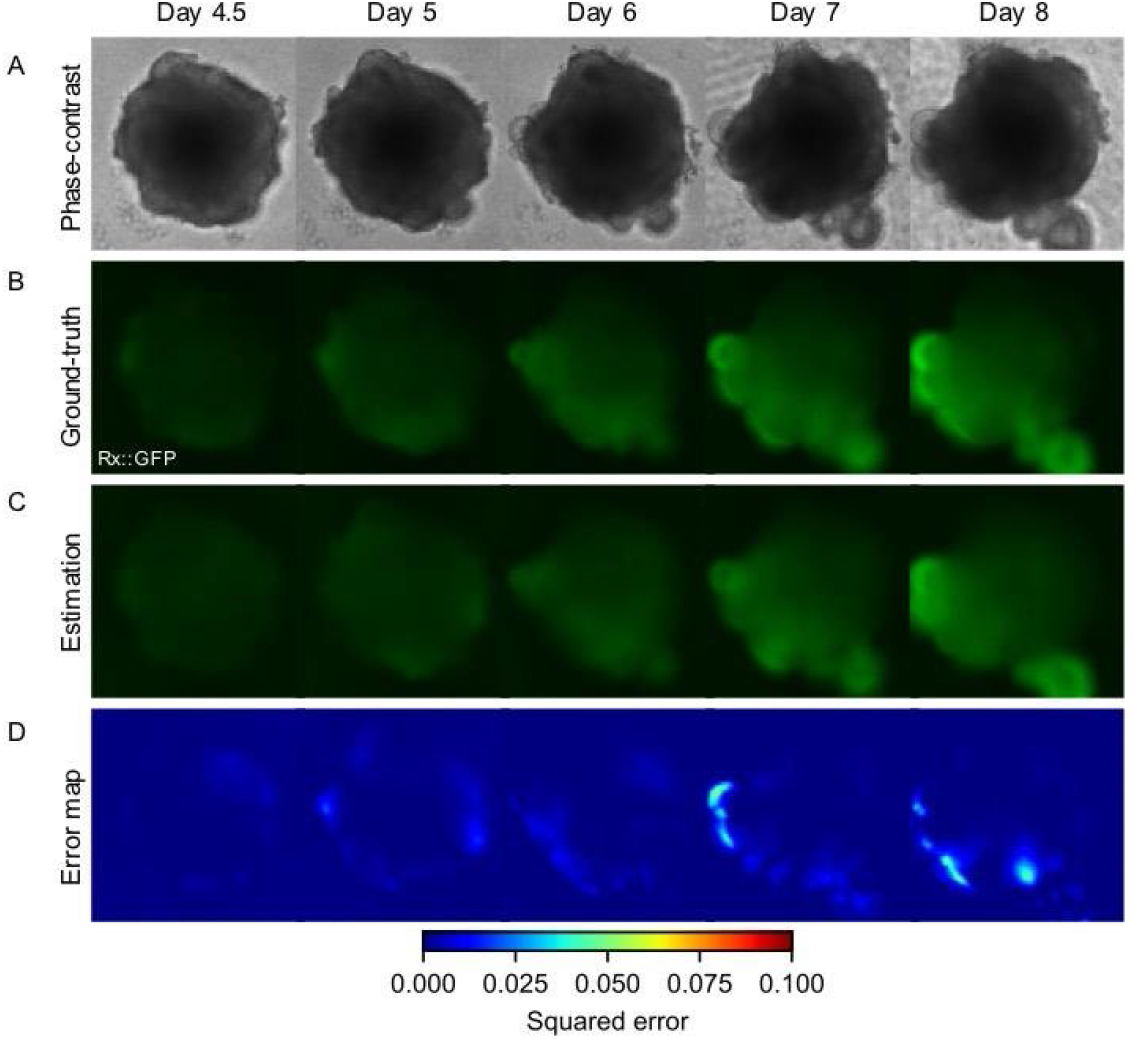
Estimated spatial Rx expression patterns during retinal organoid development. (A) Phase-contrast images from day 4.5 to day 8. (B) Captured fluorescent images of Rx as ground-truths. (C) Estimated fluorescent images with our model. (D) Error maps between captured and estimated images. The error metric was a squared error. The organoids in (A-D) are identical.

Our model successfully recapitulated the above features of Rx expression (Fig. 2C), i.e., the Rx intensity was relatively low and homogenous at days 4.5, 5, 6, and gradually increased around the evaginated tissue regions at days 7 and 8. Remarkably, the regions of high Rx expression were accurately estimated even in organoids with various morphologies. On the other hand, as the Rx intensity increases, especially around the evaginated tissue regions, the error of the estimated image from the ground-truth image increases with time (Fig. 2D).

To quantitatively evaluate the accuracy of the estimation, we statistically analyzed the estimation results at each stage. To clarify whether the model can estimate the Rx intensity in both samples with positive and negative Rx expression, each of the ground-truth and estimated fluorescent images was automatically classified into two categories with the coefficient of variation of the foreground pixels in a fluorescent image at day 8 (Fig. 3A). For each of these categories, the mean and coefficient of variation of the pixel values were calculated (Fig. 3B-E). In calculating these values, the phase-contrast images were binarized to obtain foreground and background masks, and then computed using only the foreground pixels and normalized to those of the background pixels.

**Figure 3.**
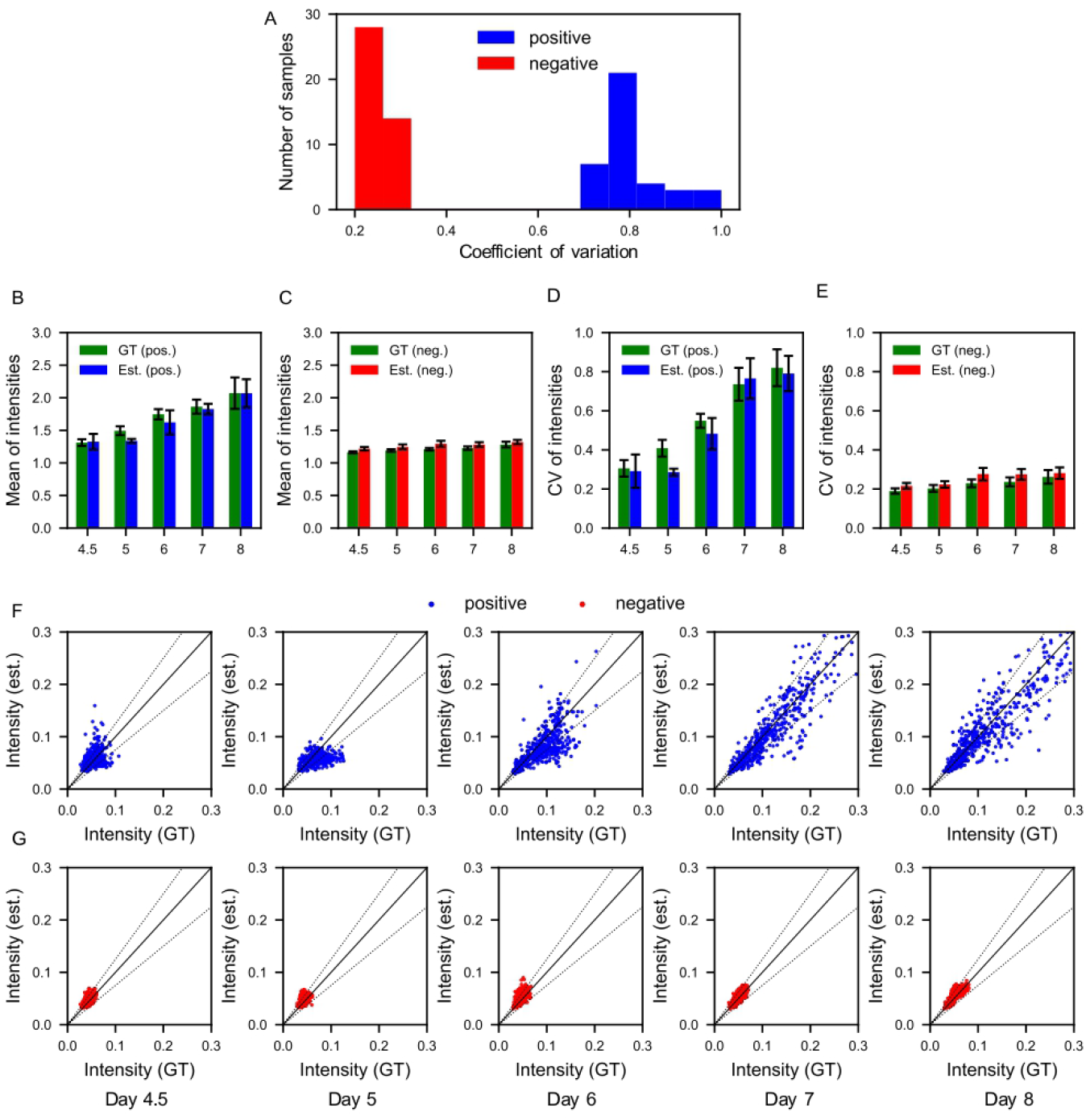
Statistical analysis of fluorescence at each developmental stage for positive and negative samples. (A) Histogram of coefficient of variation for foreground pixel values of fluorescent images at day 8. (B, C) Means of pixel values in positive and negative samples at each stage for ground-truth (green bars) and estimated fluorescent images (blue bars), respectively. (D, E) Coefficients of variation in positive samples at each stage for both ground-truth (green bars) and estimated fluorescent images (red bars), respectively. (F, G) Plots of ground-truth and estimated pixel values in positive and negative samples at each stage, respectively. Errors are 0% and 25% on the solid and dotted black lines, respectively. Error bars in (B-E) indicate standard deviations.

Positive samples showed a gradual increase in mean and intensity over the days passed (Fig. 3B). The negative sample, on the other hand, showed relatively low values from the beginning and did not change significantly over the days (Fig. 3C). Similarly, the coefficients of variation increased in the positive samples but not in the negative samples (Fig. 3D-E). These results indicate that the model successfully estimates the feature of the spatial Rx expression patterns during retinal organoid development, i.e., positive samples gradually increase Rx expressions and their heterogeneity, but negative samples do not. The intensity of the estimated images is relatively lower than the intensity of the ground-truth images in the positive samples and vice versa in the negative samples.

To clarify whether the model is capable to estimate intermediate values of the Rx expression, we analyzed the correlations between ground-truth and estimated values on foreground pixels at each stage, respectively (Fig. 3F-G). The results show that in the positive sample (Fig. 3F), the distribution of intensities is initially concentrated at low intensities and gradually expands to high intensities as the day progresses, with a wide distribution from low to high intensities. Similarly, in the negative sample, the luminance distribution is initially concentrated at low intensities, but does not expand as much as in the positive sample (Fig. 3G). These results indicate that the model successfully estimated the plausible values across all pixel intensities, demonstrating the capability of our method to infer intermediate levels of gene expression.

### Estimated spatial Rx expression patterns correspond to organoid morphologies

To determine whether the estimated Rx expression patterns correspond to tissue morphologies, we quantified the spatial distribution of Rx intensity and the mean curvature along the tissue contour around each optic vesicle (Fig. 4). For this analysis, four typical optic vesicles were selected from the positive samples and their curvature and Rx distribution were quantified. In this analysis, tissue contours were extracted and the radius of a circle passing through three points on the tissue contour was calculated as the inverse of the curvature. Moreover, the Rx intensity was measured as the average value along the depth direction from the tissue contour.

**Figure 4.**
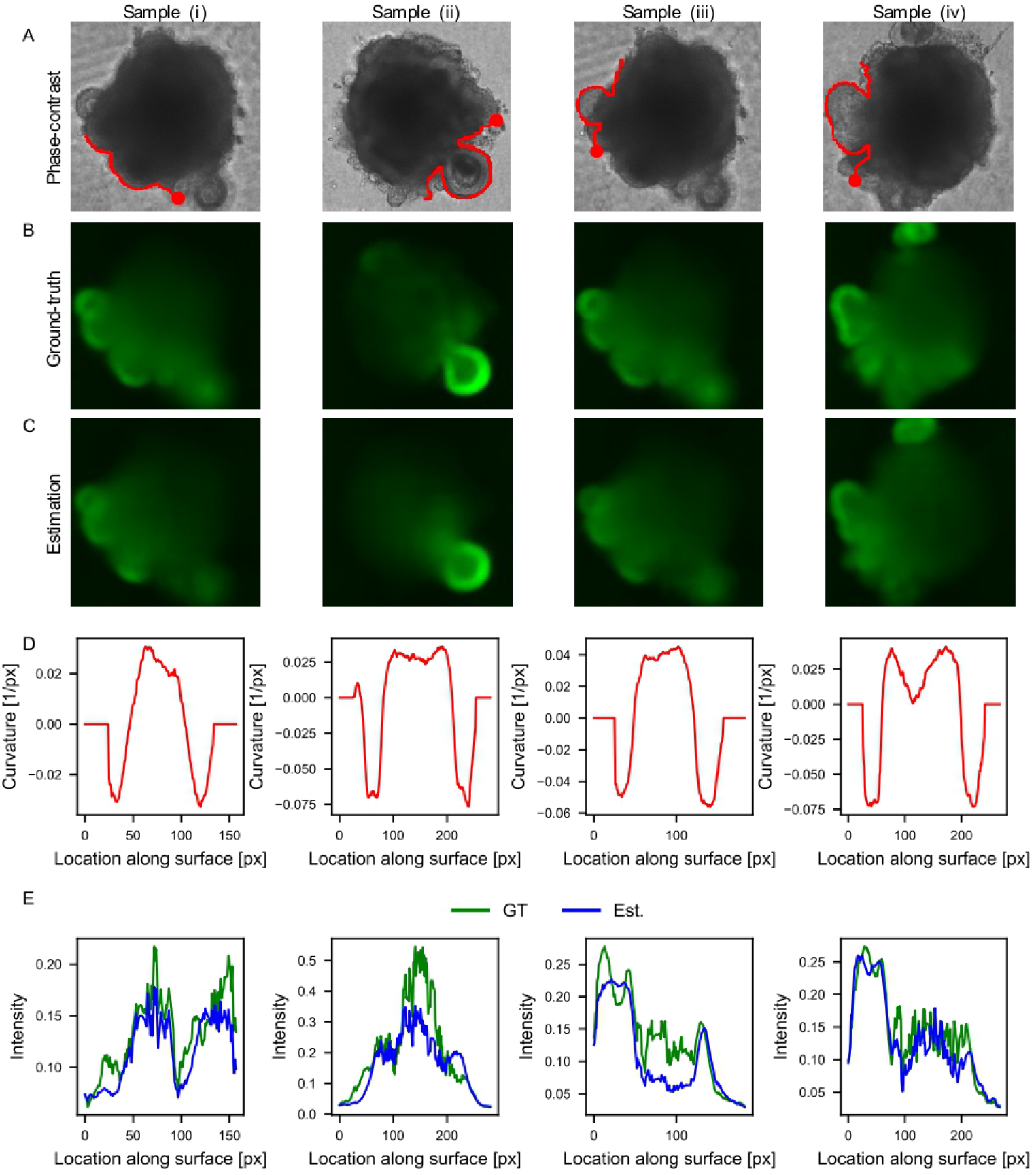
Correlation analysis of spatial Rx expression patterns and optic-vesicle morphologies. (A) Phase-contrast images. (B) Captured fluorescent images of Rx as ground-truths. (C) Estimated fluorescent images with our model. (D) Mean curvatures as a function of the distance along the organoid contour. (E) Captured and estimated fluorescent intensities of Rx along the organoid contour. The organoids in (A-C) are identical and captured on day 8. The mean curvatures and fluorescence in (D-E) are for the regions indicated by the red line starting from the red dot in (A).

Optic vesicles are hemispherical, with positive curvature at the distal portion and negative curvature at the root (Fig. 4A, D). The Rx intensity is continuously distributed around each vesicle, being highest at the distal part and gradually decreasing toward the root (Fig. 4B, E). Furthermore, because the test images were taken with a conventional fluorescence microscope, structures above and below the focal plane are included in each image. Therefore, although some images have multiple overlapping vesicles (e.g., samples iii and iv), the model successfully estimated the Rx intensity of the overlapping regions as well.

### The best balance of the training loss weights depends on the time point

MSE is commonly used as the training loss for training regression models. In addition to MSE, this model also uses cosine similarity, which can capture structural patterns from the entire image. To analyze the effect of cosine similarity on the estimation accuracy, we tested the model with different weights for both error metrics (Fig. 5). The trained models were evaluated with MSE for each test dataset on different days (Fig. 5A). The results demonstrated that cosine similarity improved the estimation accuracy at the early and intermediate stages, such as from day 4.5 to day 6. At these stages, the intensity in the differentiated region is weak, making it difficult for the network to capture structural patterns using MSE alone. Cosine similarity, on the other hand, enabled the network to learn the patterns from the weak intensity by calculating the correlation between the normalized ground-truth and the estimated images (Fig. 5B).

**Figure 5.**
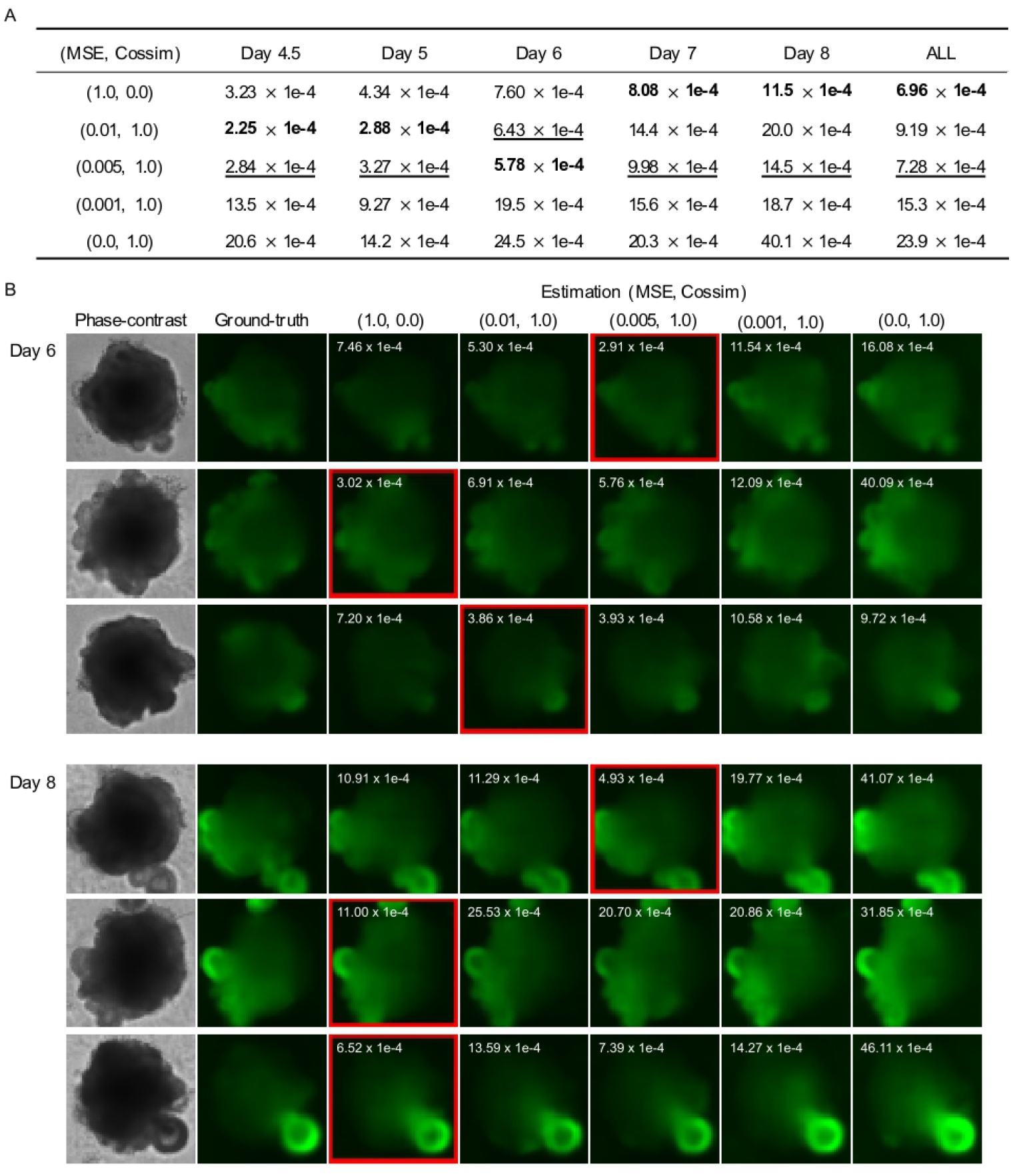
Effects of the balance of training loss on estimation accuracy. (A) Mean squared error at each stage with different hyperparameters, where bold and underlined numbers stand for the best and second best results on each day, respectively. (B) Examples of estimated fluorescent images at days 6 and 8 with different hyperparameters. The MSE of each estimated image is described in the upper left. The results with the lowest MSEs are surrounded by the red boxes.

Therefore, our model has the capability to achieve the best estimate at different stages with appropriate weight balancing.

## Discussion

To estimate spatial gene expression patterns of organoids in a non-invasive manner, we developed a machine learning-based method that converts a phase-contrast image to a fluorescent image expressing a spatial gene expression pattern. The applicability of the method was demonstrated by applying it to retinal organoids derived from mouse ESCs as an example, i.e., i) the spatial expression patterns of the retinal master gene Rx can be estimate at each developmental stage of retinal organoid development, ii) the model can be applied to both samples with positive and negative for Rx expression (retinal cs non-retinal tissues), iii) the intermediate levels of Rx expressions can be plausibly estimated, iv) the spatial Rx distributions can be estimated corresponding to organoid morphology, and v) the model can be optimized to improve the accuracy by adjusting the parameters of the training loss. These results show the potential for applying machine learning to tissues defined by Rx and its orthologous genes (Bailey et al., 2004) in various animals and at various stages.

A notable feature of this method is that it does not simply classify each organoid, instead, it estimates a spatial pattern of gene expression within each organoid. Most previous studies aim to apply image classification methods (Krizhevsky et al., 2012) to cell images, where the CNN takes a 2D image and outputs the probability of a semantic label (Kegeles et al., 2020; Kusumoto et al., 2018; Waisman et al., 2019; Zhu et al., 2021). The most relevant study to our research is the classification of retinal organoid images as differentiated or not (Kegeles et al., 2020). On the other hand, our method employs an image-to-image translation method for cell images, such as image style transfer (Isola et al., 2017), super-resolution (Ledig et al., 2017), image deburring (Dong et al., 2020), and depth (distance from a camera to an object) estimation from a monocular image (Hu et al., 2019). The image-to-image translation method outputs a 2D image and enables pixel-wise intensity estimation. Therefore, this method is novel in that it can quantitatively estimate the spatial pattern of gene expression in each organoid.

This method can automatically and non-invasively estimate the spatial patterns of specific gene expressions in each organoid. This feature can assist in the evaluation of tissues in a wide range of biological and basic medical research. For example, this method can infer specific tissue regions without genomic alterations. Therefore, it is possible to fully analyze the endogenous gene network while maintaining the activation of the two endogenous alleles. Moreover, it avoids the ethical issues that prevent the use of genomic alterations for cell, tissue, and organ transplantation. In addition, the high throughput and automation of this method greatly reduce the time and effort required for genome editing. In addition, the method does not require human judgment, allowing for objective and robust evaluations.

In this study, the developed method was applied to Rx expression in retinal organoids as an example, while it is likely to be applicable to other organoids and other genes. In addition, the method may be applicable not only to organoids, but also to embryonic tissues, pathological tissues, and other biological tissues in principle. However, this method may not be applicable to all genes because gene expressions do not always coincide with tissue structures. Moreover, even for genes that are applicable, the model needs to be trained each time depending on the tissue type, sample preparation, and imaging method, while the estimation accuracy can be improved by adjusting the parameters according to the experimental situation (Fig. 5). In addition, while the images used for training in this study were captured with a regular fluorescence microscope, the estimation accuracy could be further improved if clearer images could be obtained with a confocal microscope or other means. Thus, our approach opens a new avenue to assess the genetic activities of cells in organoids and other tissues in the wide range of biology and regenerative medicine.

## Materials and methods

### Mouse ES Rx-reporter cell line and 3D culture

Mouse ES cells (EB5, Rx-GFP) were maintained as described in the previous study (Eiraku et al., 2011; Watanabe et al., 2005). The cell line that we used is a subline of the mouse embryonic stem cell line, EB5 (129/Ola), in which the GFP gene was knocked-in under the Rx promoter. It is available from RIKEN BioResource Research Center (ID: AES0145). The SFEBq culture method was performed as described in the previous study (Eiraku et al., 2011). In this culture, 3,000 cells were suspended and 3.5% Matrigel (Corning®) was added to each well of a 96-well plate to form cell aggregates.

### Automated imaging

Both phase-contrast and fluorescent images were captured using a Keyence All-In-One Fluorescence Microscope BZ-X800. A 10X objective lens was used. On each day from 4.5 to 8 during SFEBq culture, 14 images were taken for each organoid at different Z positions with 15 µm intervals within 195µm. The resolution of both phase-contrast and fluorescent images is 960 × 720, from which the 512 × 512 regions where organoids appear were extracted; phase-contrast images were binarized and the median pixels in the vertical and horizontal axes were computed, and then the 512 × 512 regions centered at these median pixels were cropped.

### Quantification of curvature and Rx distributions

To quantify tissue curvature and Rx distributions, tissue contours were first manually extracted from the phase-contrast images. Then, the radius of a circle passing through three points on the tissue contour (the point where the curvature is measured and two points 81.6 µm before and after that point) was calculated and its reciprocal was defined as the curvature. The average of the intensity from the point at which the Rx expression intensity is measured on the tissue contour to 13.6 µm vertically from the tangent line of that point was also measured.

### Training loss

The training loss ℒ is computed from the output fluorescent images and the corresponding ground-truth images. The trainable parameters *θ* (the weights of these convolution kernels and biases for CNNs) are optimized by minimizing the training loss:

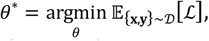

where 𝒟 is the training dataset, and **x** and **y** are a phase-contrast and the corresponding fluorescent image, respectively. A naïve loss function is the mean squared error (MSE), where the training loss is computed pixel by pixel as follows:

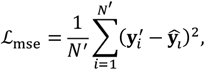

where 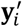 and 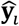 are the values at the *i*-th pixel in the ground-truth and estimated images, respectively. *N*′ is a number of pixels. In this study, we additionally use the cosine similarity as follows:

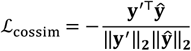

This makes the network learn a global structure, because the loss is computed considering the whole pixels simultaneously. We combine the above two losses as follows:

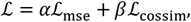

where *α* and *β* are hyperparameters to control the contribution of each loss.

### Implementation details

Our network was implemented in PyTorch. Training was performed on an NVIDIA A100 GPU. The size of a mini-batch was 16 for the training loss. The encoder was initialized by the ResNet50 pre-trained on the ImageNet (Deng et al., 2009). The optimizer was Adam (Kingma and Ba, 2015) with the default hyperparameters (*lr* = 10^−3^, *β*_1_ = 0.9, and *β*_2_ = 0.999). The learning rate was multiplied by 0.1 at every 20 epochs. The network was trained for 50 epochs and we adopted the best model on the validation dataset.

## Author contributions

Y.F.: Conceptualization; Methodology; Software; Writing – original draft; Writing – review & editing. I.S: Data curation; Investigation. I.S.: Data curation; Investigation. N.T: Writing – review & editing. A.H.: Conceptualization; Methodology; Writing – review & editing. T.F.: Conceptualization; Methodology.; Writing – review & editing. S.O.: Conceptualization; Funding acquisition; Project administration; Resources; Writing – original draft; Writing – review & editing.

## Acknowledgements

This work was supported by the Japan Science and Technology Agency (JST), CREST Grant No. JPMJCR1921 (S.O.) and PRESTO Grant No. JPMJPR2025 (T.F.); the Japan Agency for Medical Research and Development (AMED), Grant No. 21bm0704065h0002 (S.O.); the Japan Society for the Promotion of Science (JSPS), KAKENHI Grants No. 21H01209, 22K18749, and 22H05170 (S.O.); and the World Premier International Research Center Initiative, Ministry of Education, Culture, Sports, Science and Technology (MEXT), Japan (S.O).

